# Microbial Engraftment and Efficacy of Fecal Microbiota Transplant for *Clostridium difficile* Patients With and Without IBD

**DOI:** 10.1101/267492

**Authors:** Robert P Hirten, Ari Grinspan, Shih-Chen Fu, Yuying Luo, Mayte Suarez-Farinas, John Rowland, Eduardo J. Contijoch, Ilaria Mogno, Nancy Yang, Tramy Luong, Philippe R. Labrias, Inga Peter, Judy H. Cho, Bruce E. Sands, Jean Frederic Colombel, Jeremiah J. Faith, Jose C. Clemente

## Abstract

**Background & Aims:** Recurrent and refractory *Clostridium difficile* infections (CDI) are effectively treated with fecal microbiota transplant (FMT). Uncertainty exists regarding the effectiveness of FMT for CDI with underlying inflammatory bowel disease (IBD), its effects on disease activity and its effectiveness transferring the donor microbiome to patients with and without IBD. This study aims to determine FMTs effectiveness in subjects with and without IBD, its impact on IBD activity, the level of microbiome engraftment, and predictors of CDI recurrence.

**Methods:** Subjects with and without IBD who underwent FMT for recurrent or refractory CDI between 2013 and 2016 at The Mount Sinai Hospital were followed for up to 6 months. The primary outcome was CDI recurrence 6 months after FMT. Secondary outcomes were (1) CDI recurrence 2 months after FMT; (2) Frequency of IBD flare after FMT; (3) Microbiome engraftment after FMT; (4) Predictors of CDI recurrence.

**Results:** Overall, 134 patients, 46 with IBD, were treated with FMT. There was no difference in recurrence in patients with and without IBD at 2 months (22.5% vs 17.9%; p=0.63) and 6 months (38.7% vs 36.5%; p>0.99). Proton pump inhibitor use, severe CDI, and comorbid conditions were predictors of recurrence. The pre-FMT microbiome was not predictive of CDI recurrence. Subjects with active disease requiring medication escalation had reduced engraftment. There was no difference in engraftment based on IBD endoscopic severity at FMT.

**Conclusions:** IBD did not affect CDI recurrence rates 6 months after FMT. Pre-FMT microbiome was not predictive of recurrence, and microbial engraftment was dependent on IBD treatment escalation but not on underlying disease severity.

## INTRODUCTION

*Clostridium difficile* infection (CDI) is one of the most common health-care associated infections and is associated with significant morbidity and mortality.^1^ After initial antibiotic therapy, 10 to 20% of patients will experience a recurrence, and up to 65% will recur after subsequent episodes.^2,3^ Generally, the first recurrence is treated with the same antibiotic regimen used for the initial infection, while a prolonged vancomycin course and fecal microbiota transplant (FMT) are used for the second and third recurrences, respectively^4^.

CDI has been associated with alterations of the intestinal microbiome, generally reducing bacterial diversity and the abundance of Bacteroidetes and Firmicutes phyla.^5^ FMT effectively treats recurrent CDI in approximately 90% of patients by replacing the patient’s aberrant microbiome with a donor-like microbiome. While FMT is currently used in clinical practice to treat CDI in patients with IBD, studies have demonstrated variable efficacy in this population.^6,7^ There are concerns regarding the use of FMT in patients with underlying IBD due to the frequent use of concomitant immunosuppressive agents and the possibility of worsening IBD activity. Several studies found a worsening of IBD activity in approximately 23% of patients post FMT.^6–9^ Furthermore, it is unknown whether microbiome engraftment is lower in patients with concomitant IBD compared to those with CDI only, which could result in increased recurrence rates.

Given these questions, the goal of this study is to determine its long-term effectiveness in the treatment of CDI and the predictors of post-FMT recurrence in patients with and without IBD. Evaluation of the microbiome was also performed in a subset of patients to assess the impact of IBD on engraftment and its subsequent risk of relapse.

## MATERIALS AND METHODS

### Study Design

This is a longitudinal retrospective cohort study including all patients with and without IBD who underwent FMT for recurrent or refractory CDI between 2013 and 2016 at The Mount Sinai Hospital (New York, United States). The study was approved by the Mount Sinai Institutional Review Board. Eligibility criteria were for recurrent CDI characterized as either at least 3 episodes of mild to moderate CDI and failure of a 6 to 8-week taper with vancomycin, or at least 2 episodes of severe CDI resulting in hospitalizations and associated with significant morbidity, and for refractory CDI including either persistent moderate to severe CDI not responding to standard therapy (vancomycin) for at least 1 week, or severe (including fulminant CDI) with no response to standard therapy after 48 hours.

FMT was obtained from healthy donors, CIPAC Therapeutics or OpenBiome, with healthy donors screened as previously published.^10^ (see Suppl. Materials). Baseline demographic data as well as the medical and surgical history for all patients were collected. IBD activity at the time of FMT and at 2 and 6 months post-transplant was characterized utilizing clinical disease activity scores (Harvey-Bradshaw index [HBI] for CD and the partial Mayo Score for UC). Endoscopic IBD severity was captured at the time of FMT utilizing endoscopic grading systems (Simple Endoscopic Score for Crohn’s Disease [SES-CD] and The Mayo endoscopic subscore). IBD related medications were captured prior to FMT, and longitudinally in patients in whom the microbiome was analyzed to assess for therapeutic escalation. Escalation was defined as the need to initiate new IBD treatment, including corticosteroids, or the need to change the current medication. The severity of the CDI was defined by the 2013 American College of Gastroenterology guidelines^3^.

The primary outcome was late CDI recurrence at 6 months after initial FMT in patients with and without IBD. The secondary outcomes were (1) Early CDI recurrence at 2 months after initial FMT; (2) Frequency of IBD flare at 2 and 6 months after initial FMT; (3) Microbiome engraftment after FMT; (4) Predictors of CDI recurrence after initial FMT. Successful FMT was defined as a resolution of diarrhea within 8 weeks of FMT and no need for re-initiation of therapy. Recurrence of CDI is defined as a recurrence of diarrhea and laboratory confirmation of *C. difficile* in the stool. An IBD flare was diagnosed by the physician treating the patient.

### Microbiome Data Generation and Analysis

A subset of 18 subjects with (n=9) and without (n=9) IBD were analyzed before and up to 12 months after initial FMT. Samples for microbiome analysis were collected prior to FMT, at the time of FMT, within 48 hours after transplant, 1 week after FMT, 4 weeks after FMT, 8 weeks after FMT, 6 months after FMT and 12 months after FMT. These 18 subjects received fresh FMT from one of 11 healthy donors, who also had their stool analyzed. Fecal microbiota was analyzed utilizing 16S rRNA sequencing as described previously.^11^ (see Suppl. Materials). Metagenomic function was predicted using PICRUSt and differential analysis of pathways was performed using STAMP^12,13^.

### Statistical Analysis

Baseline comparisons of categorical data in subjects with and without IBD were conducted using Fischer’s exact test and the Chi Square test. The t-test was used for continuous data. Recurrence rates of CDI in IBD and non-IBD groups were presented with 95% confidence intervals, computed using the results of the proportion test within each group, and compared between the groups at 2 and 6-month endpoints using Fischer’s exact test. The time to first CDI recurrence was compared using the Log-rank test and presented as a Kaplan-Meier curve. Changes in continuous outcomes over time were compared between IBD and non-IBD groups using Linear Mixed-Effects Models. To evaluate clinical variables as predictors of CDI recurrence a two-step strategy was set up where the most robust predictors were identified by combining multiple imputations and regularized regression techniques and fit to a multivariable regression model.

## RESULTS

### Patient Population

A total of 134 patients with CDI were treated with FMT, of which 46 had underlying IBD (Suppl. Table 1). Among IBD patients, 27 patients had ulcerative colitis (UC), 18 patients had Crohn’s disease (CD) and 1 patient had indeterminate colitis (Suppl. Table 2). 64% of the cohort were women and the average age was 53 years. The cohort with IBD was significantly younger than the non-IBD cohort (mean age 38.8 versus 60.3 years, p<0.001). The indication for FMT was for recurrent CDI in 89 patients and refractory CDI in 44 patients. This did not differ between the IBD and non-IBD groups (p=0.39). 21.6% of FMTs were performed in the inpatient setting and 78.4% in the outpatient setting, with fresh and frozen stool used in 51.5% and 48.5% of FMTs respectively. 51.5% of patients were hospitalized within the 90 days prior to FMT which was significantly more frequent in non-IBD patients compared to IBD patients (58% versus 39.1%; p=0.04), as was the percentage of patients requiring a past hospitalization for CDI (55.7% versus 34.8%; p=0.02). At the time of fecal transplant 91.3% of IBD patients were receiving an immunosuppressive agent. At FMT, 37 (82%) patients with IBD had evidence of endoscopic disease activity. 18.8% of the cohort had a severe CDI, with severity not differing between those with and without IBD (p=0.22).

Thirteen subjects (9.7%) underwent a previous FMT in the past, 6 (13%) in the IBD cohort and 7 (8%) in the non-IBD cohort (p=0.37). We noted differences in comorbid conditions between the two groups, with significantly less patients with IBD having hypertension (p<0.001), cardiovascular disease (p=0.006), diabetes mellitus (p=0.01), and liver disease (p=0.03). Additionally, patients with IBD were less likely than patients without IBD to have diverticulosis seen on colonoscopy at the time of FMT (4.3% versus 40.2%; p<0.001).

### CDI Outcomes and Predictors of Failure

23 out of 118 (19.5%) patients with follow up at 2 months and 31 out of 83 (37.3%) patients with follow up at 6 months suffered from recurrent CDI after the initial FMT. Subjects with IBD did not have a higher rate of CDI recurrence at 2 months (22.5% versus 17.9%; p=0.63) or 6 months (38.7% versus 36.5%; p>0.99) compared to the non-IBD group (Figure 1). The rate of recurrence was equivalent between the IBD and non-IBD group at both 2 months (p=0.03) and 6 months (p=0.02). Similarly, there was no difference between the groups in time to first CDI recurrence (p=0.46). 18 of the 107 subjects (16.8%) with available follow up data required repeat FMT by 2 months of follow up, which was not significantly higher in the group with IBD compared to those without IBD (17.2% versus 16.7%; p>0.99). At 6 months, 21 out of the 76 patients (27.6%) in whom data was available required repeat FMT, which was not significantly higher in the IBD group than without IBD (25.0% versus 29.2%; p=0.70). In the 6 month follow up, there were no serious adverse events noted secondary to FMT. At 6 months, there was no difference between colectomy rates in the IBD and non-IBD groups (12.9% versus 9.5%; p=0.72).

**FIGURE 1.**
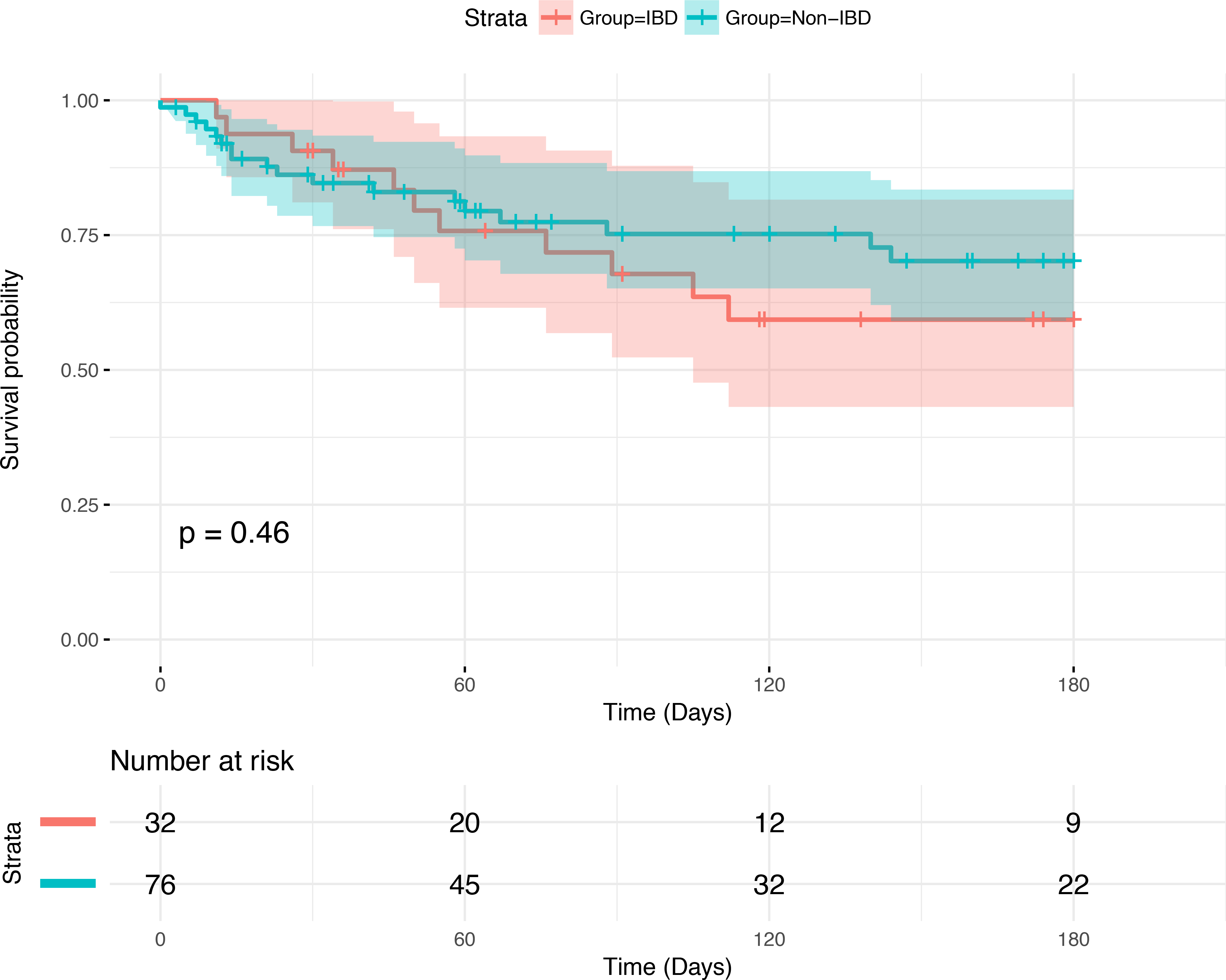
Survival analysis for time to first CDI recurrence from the date of initial FMT with Log-Rank test results. Censored at 6 months.

Univariate analysis did not reveal any factors associated with the risk of CDI recurrence at 2 months. CDI recurrence at 6 months was associated with the use of proton pump inhibitors (p=0.01), FMT performed as an inpatient (p=0.02), and a lower hemoglobin (p=0.02) (Suppl. Table 3). At 2 months and 6 months, respectively, IBD type (p=0.71, p=0.13), immunosuppression at FMT (p=0.55, p>0.99), and IBD severity at FMT (p>0.99, p=0.63) were not predictors of CDI recurrence. Based on the final logistic regression model, proton pump inhibitor use (p=0.045), severe CDI at the time of FMT (p=0.005), and hypertension (p=0.03) were all associated with an increased risk of CDI recurrence at 6 months.

### IBD Related Activity Over Time

Overall, 6 out of 37 (16.2%) and 15 out of 27 (55.6%) subjects with follow up at 2 and 6 months post FMT, respectively, had an IBD flare. A linear mixed effects model was used to calculate the least-squares means (LSM) of the HBI and Partial Mayo score over time. There was not a significant change in HBI scores over time (p=0.84) when comparing baseline (LSM 6.6; CI 4.3-8.9), 2 month (LSM 6.8; CI 4.2-9.4) and 6 month (LSM 6.1; CI 3.4-8.8) scores. Partial Mayo scores did not significantly change over time (p=0.18) between baseline (LSM 3.9; CI 3.1-4.6), 2 month (LSM 3.2; CI2.3-4.1) and 6 month (LSM 3.2; CI 2.2-4.1) values.

### Microbiome of CDI patients pre-FMT

The microbiome of patients before FMT was significantly different than their donors, with a lower alpha diversity (Figure 2A), distinct beta diversity (Figure 2B; PERMANOVA p=0.02), and depletion in Bacteroides, Lachnospiraceae, and Faecalibacterium (Supplementary Figure 1). There were no significant differences in alpha diversity (Figure 2C; p=0.31) or beta diversity (Figure 2D; p=0.45) between patients with and without IBD pre-FMT.

**FIGURE 2.**
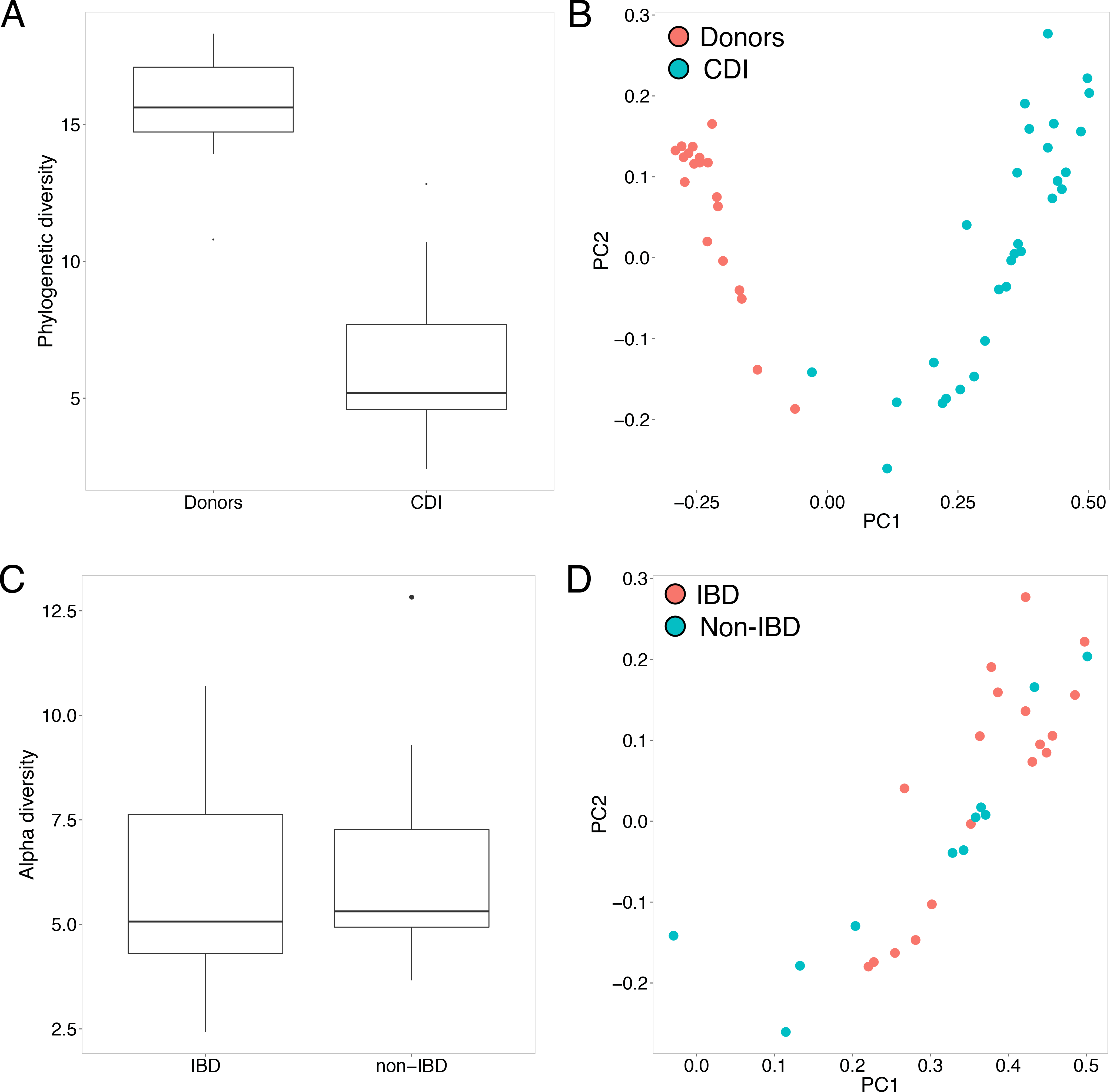
(A) Alpha diversity of donors and CDI patients pre-FMT (B) PCoA plot based on beta diversity distances of donors and CDI patients pre-FMT (C) Alpha diversity of CDI patients pre-FMT with and without underlying IBD (D) PCoA plot based on beta diversity distances of CDI patients pre-FMT with and without underlying IBD.

### FMT Induces Significant Changes in Microbiome Composition and Diversity

Bacterial alpha diversity increased significantly from 6.3 ± 2.4 before transplant to 13.4 ± 3.5 immediately after, and remained high throughout the 12 month follow up period (Figure 3A). No significant differences were observed between the overall alpha diversity of donors and patients post-FMT. Beta diversity was also significantly distinct before and after FMT (Figure 3B; PERMANOVA *p*=0.001), with a decrease in microbiome distance between recipients and donors immediately after transplant and maintained for the duration of the study (Figure 3C). Principal coordinate analysis (PCoA) revealed a gradient along the first principal coordinate with time since transplant (Figure 3D; R^2^=0.501, p=1.42e-15). These changes were mostly mediated by a significant enrichment in Bacteroides, Lachnospiraceae, Faecalibacterium, Blautia, and Ruminococcaceae after FMT (Figure 3E). Finally, a random forest classifier trained on microbiome composition accurately predicted whether samples from the patients were obtained before or after FMT (Supplementary Figure 2).

**FIGURE 3.**
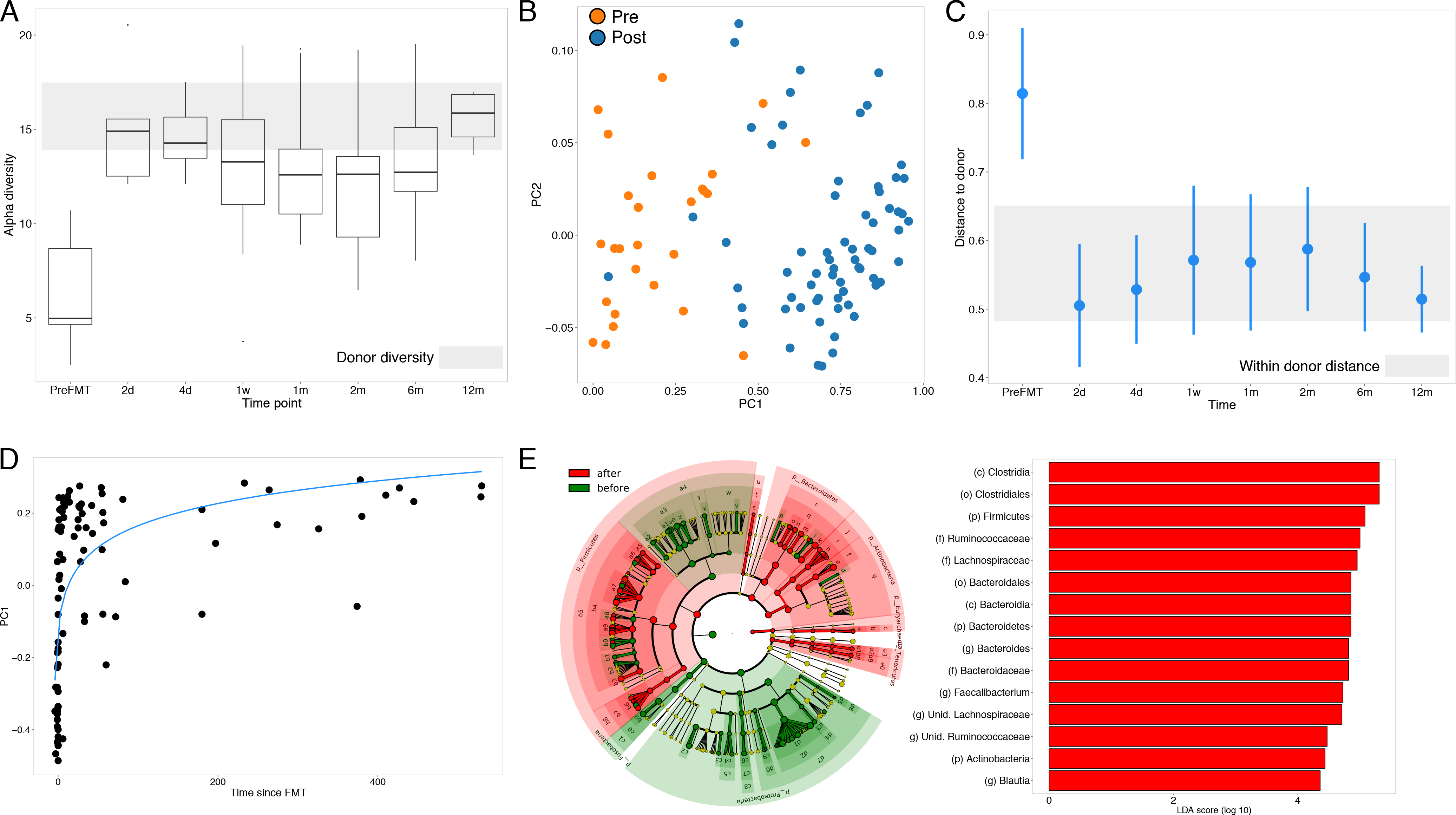
(A) Longitudinal alpha diversity over time in CDI patients. (B) LEfSe analysis of patients before and after FMT (C) PCoA plot based on beta diversity distances before and after FMT. (D) First principal coordinate versus time since transplant. Curve represents log_10_ fit. (E) LEfSe analysis comparing microbial composition before (green) and after (red) FMT. Represented are all taxa significantly distinct with LDA scores;>;2.0.

### Bacterial Engraftment after FMT is Associated with Changes in IBD Treatment

Although FMT results in significant changes in the microbiome of patients, we did not observe significant differences between the microbiome of patients with (IBD) and without IBD (Non-IBD) post-FMT (Supplementary Figure 3). Stratification of the IBD group based on disease activity, comparing those with mild endoscopic disease (n=2) against those with moderate to severe endoscopic disease (n=7), revealed no significant differences in the microbiome. Importantly, the IBD patients who required a change in IBD-related medication after FMT (IBD^e^ n=6) exhibited a blunted increase in bacterial diversity immediately after FMT compared to those who did not require change in medications post FMT (IBD^s^ n=3) (Figure 4A). Beta diversity was also altered in the IBD^e^ group with significant differences between this group and all others (Figure 4B). These changes immediately after FMT persist over time: alpha diversity of the IBD^e^ patients failed to reach levels observed in the donors, while the non-IBD/IBD^s^ groups were within diversity levels of healthy donors (Figure 5A). Beta diversity also exhibited a distinct temporal pattern in the IBD^e^ group (Figure 5B). Network analysis also revealed differences in microbiome structure after FMT, with non-IBD/IBD^s^ patients being enriched in clusters of Bacteroides, Clostridiales, and Faecalibacterium, while IBD^e^ were enriched in Enterobacteria and Lactobacillus (Supplementary Figure 4). Furthermore, the non-IBD/IBD^s^ network had lower clustering coefficient and centralization than the IBD^e^ network (0.303 and 0.155 vs 0.421 and 0.168), while exhibiting a longer average path length (3.78 vs 3.36).

**FIGURE 4.**
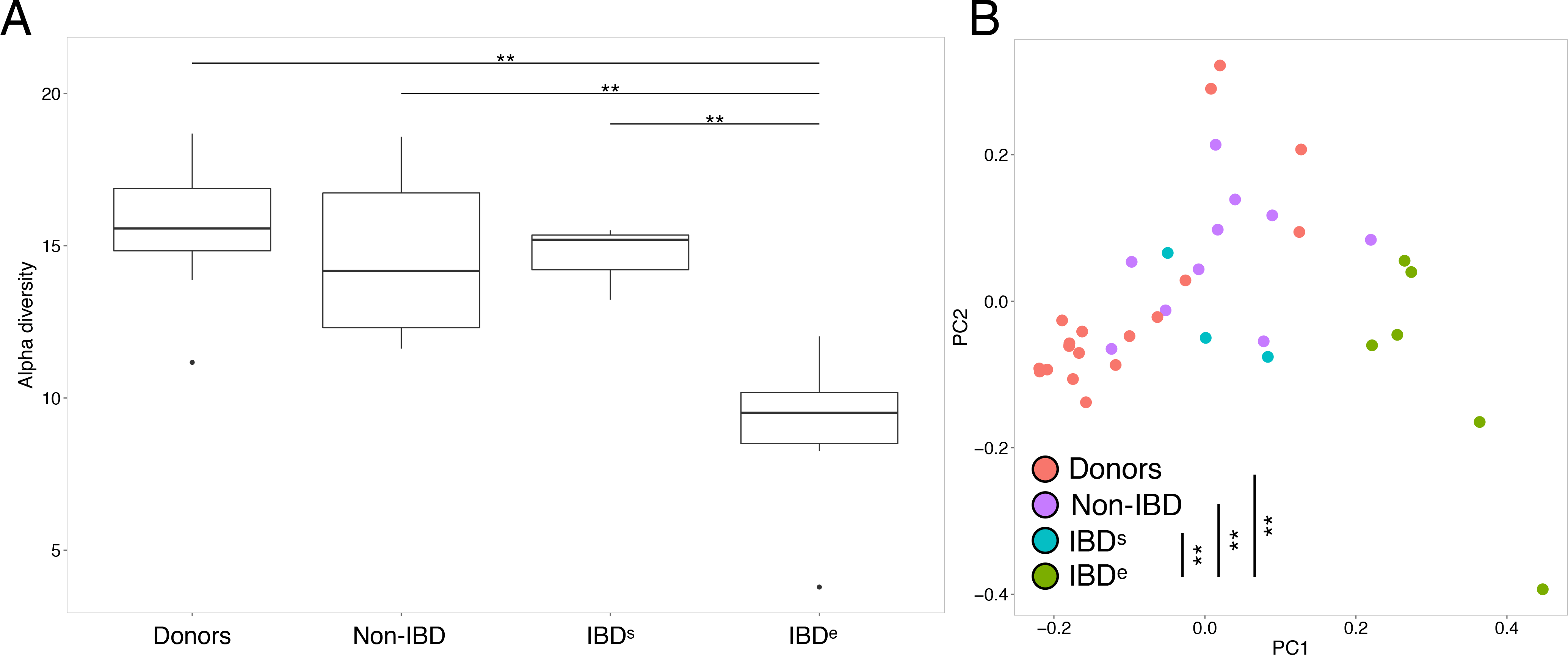
(A) Alpha diversity in IBD^+,e^ versus IBD^+,s^ and IBD^−^ group immediately after FMT (B) PCoA plot based on beta diversity distance immediately after FMT in IBD^e^ versus IBD^s^ and non-IBD groups.

**FIGURE 5.**
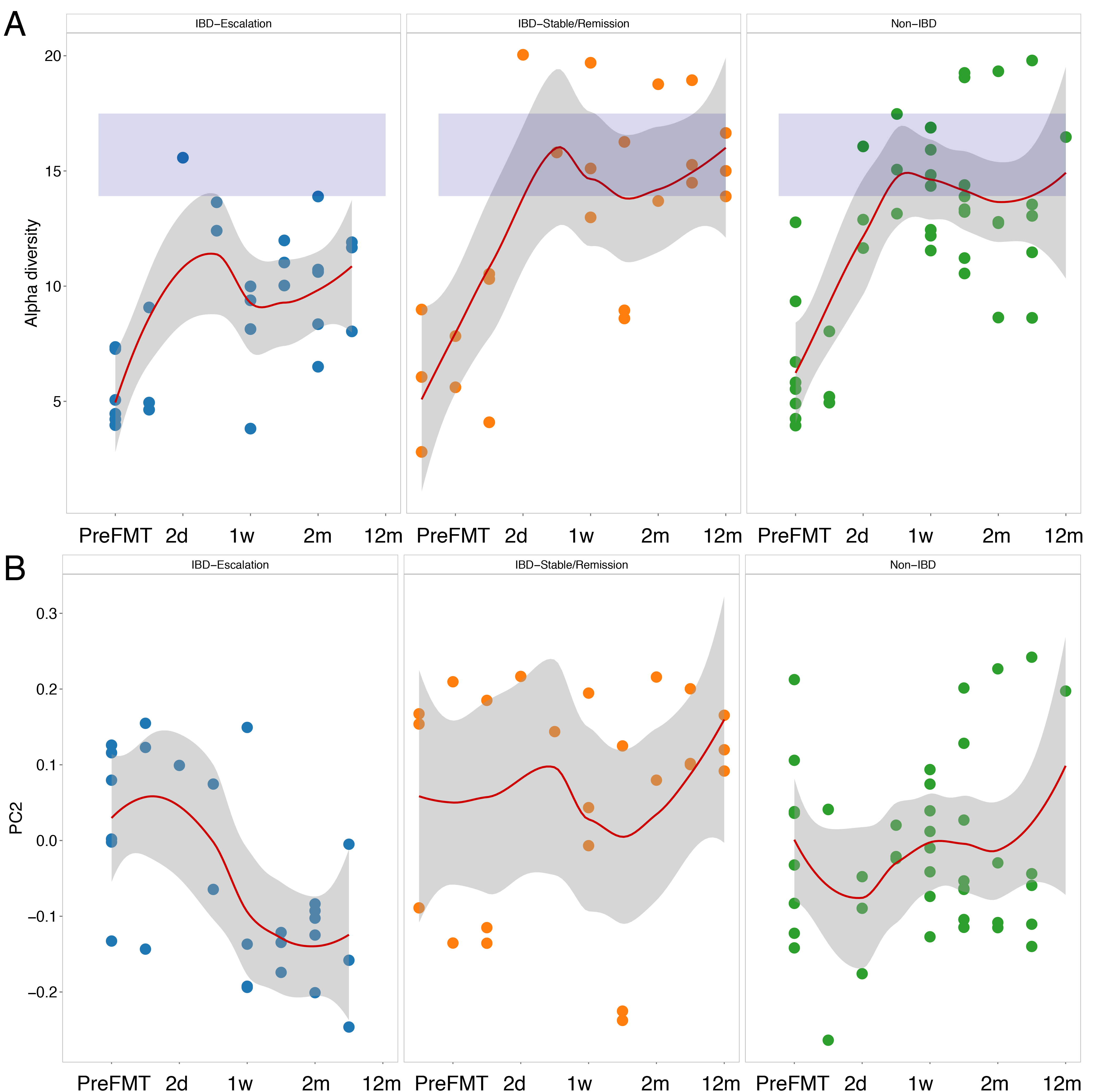
(A) Alpha diversity in IBD^e^ versus IBD^s^ and non-IBD group immediately after FMT. (B) Changes in second principal coordinate over time in the IBD^e^ group versus IBD^s^ and non-IBD. Curves smoothed over time using local regression, with gray area representing the confidence interval.

### IBD Therapy Escalation is Associated with Functional Shifts in the Microbiome

Bacterial functions were also significantly different between IBD/IBD^s^ and IBD^e^ groups. We observed a decrease in pathways associated with non-sulfur containing amino acids (lysine biosynthesis, histidine metabolism), enrichment in bacterial homeostasis during oxidative stress (glutathione metabolism), and enrichment in clinical disease activity (lipopolysaccharide biosynthesis) (Figure 6). Overall, these results suggest an enrichment of functions associated with pathogenicity in the IBD^e^ cohort.

**FIGURE 6.**
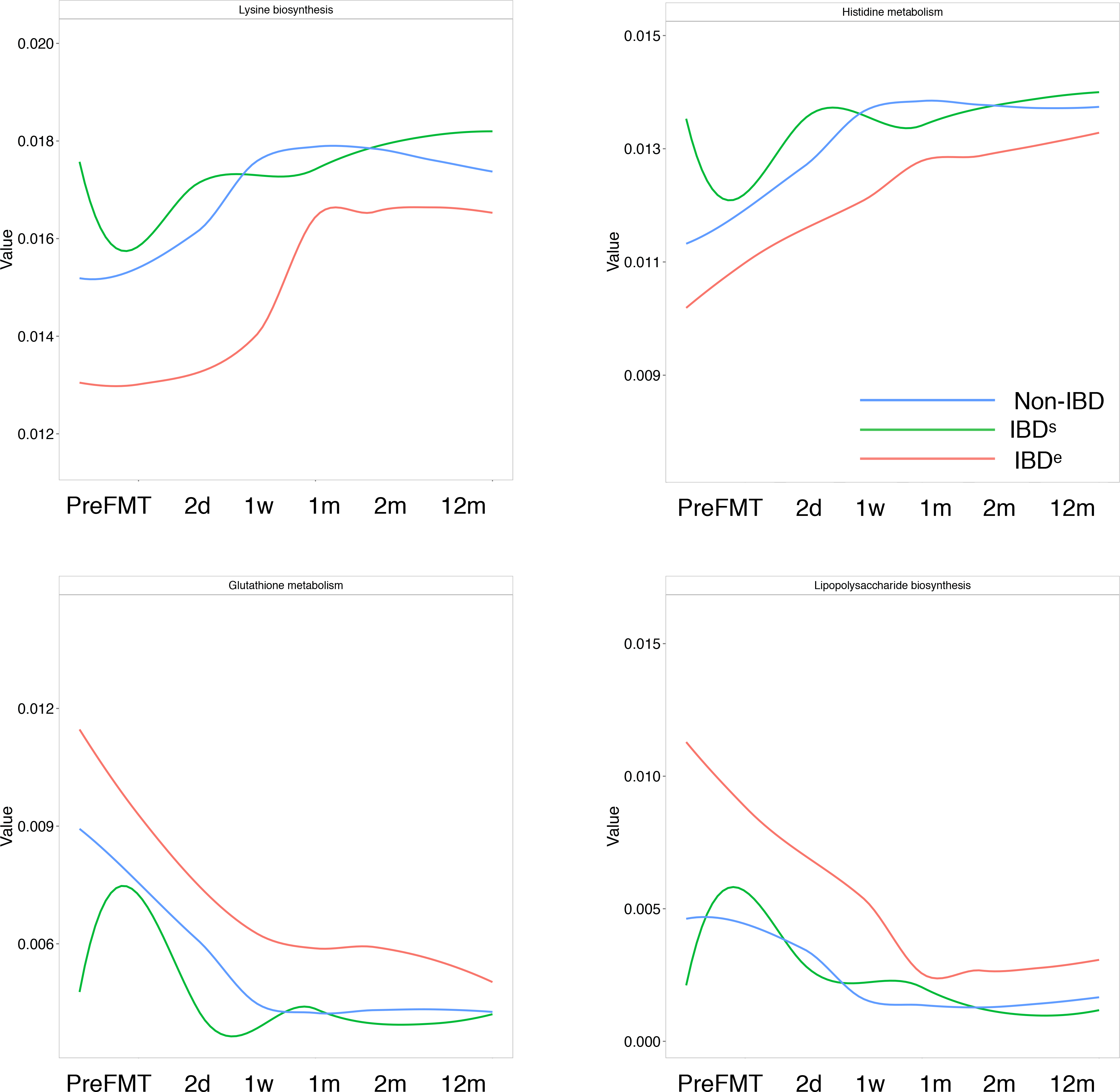
PICRUSt-predicted bacterial pathways differentially enriched in IBD^e^ (red) versus IBD^s^ (green) and non-IBD (blue). Curves smoothed over time using local regression.

## DISCUSSION

We report the first study combining a long-term evaluation of the microbiome and the risk of recurrent CDI up to 6 months after FMT in a cohort of subjects with and without IBD. Our results support two important conclusions: First, IBD does not significantly increase the risk of recurrent CDI after FMT; second, microbiome engraftment after fecal transplant is not influenced by the presence or absence of underlying IBD or the degree of disease activity, but rather escalation of IBD therapy.

The vast majority of FMT studies to date have focused on efficacy outcomes at 1-3 months post-FMT, with few long-term studies published in the literature.^6,7^ Many studies exclude subjects with severe CDI, a known predictor of CDI recurrence, which may explain the lower success rate of FMT observed in our cohort compared to others.^6^ However, our results indicate that there is a non-trivial late recurrence that occurs between 2 and 6 months (18.5% versus 37.3% recurrence) post FMT. Underlying IBD has been proposed as a risk factor for late recurrence of CDI,^14^ but in our cohort we found no difference in recurrent CDI at 6 months between IBD and non-IBD patients. Identified risk factors for CDI recurrence after FMT have included severe CDI, inpatient status, the number of previous CDIs and a low albumin at the time of FMT.^7,15^ These predictors were for short term relapse, generally within 2 months of fecal transplant. We identified long term predictors of relapse including severe CDI, proton pump inhibitor use, and the comorbid condition hypertension. Low albumin and an inpatient location of FMT were not found to be predictive of FMT failure in our study; however, these reflect CDI severity, which we found to be predictive of recurrence.

There is continued controversy regarding the impact of IBD on the efficacy of FMT. Khoruts and colleagues had demonstrated a negative effect of IBD on the success of FMT, with 2 month CDI clearance rates of 74% versus 92.1% in those with and without IBD, while Fischer and colleagues failed to identify IBD as a predictor of early failure.^6,7^ To address these discrepancies our primary outcome was long term CDI recurrence, which provided an extended follow up period to capture possible relapses that might be missed with shorter durations of follow up. At 2 and 6 months post-FMT, underlying IBD was not found to influence recurrence and was not found to be a predictor of relapse. Additionally, we did not find IBD type or severity to be predictive of recurrence.

It has been hypothesized that a deficient immune response in subjects with IBD impact the microbiome, explaining the reduced efficacy of FMT in subjects with IBD that is observed in some studies.^6,16,17^ We longitudinally analyzed the microbiome in a subset of our patients to assess the impact of IBD on engraftment and its change over time. While others have described a reduced increase in diversity in patients with concomitant IBD compared to those without^18^, we observed no such difference either before or after FMT. These findings support the clinical outcomes we observed, underscoring that IBD status does not necessarily impact the efficacy of FMT.

There is concern that the use of FMT to treat CDI can provoke a flare or worsening of underlying IBD activity. A recent meta-analysis found that the risk of an IBD flare after FMT is as high as 22.7%.^8^ We found that 16% of subjects developed a flare of their IBD within 2 months of their FMT, which is in line with previous reports from another large series.^9^ When examining the HBI and partial Mayo scores of the larger IBD cohort, we found no significant increase in IBD activity at 2 and 6 months post-FMT, showing that most patients tolerate FMT without an appreciable worsening of disease activity. This further reflects that those subjects with a flare post-FMT are generally able to have their disease brought under control. While providers should be aware that the risk of disease flare after FMT exists, our findings support a relatively stable disease course over time.

FMT results in rapid changes in the microbiome of patients after transplantation. Our analysis did not reveal any difference in diversity between subjects who did and did not recur after FMT, which is consistent with the findings of others.^19^ Subgroup analysis of the microbiome did not find differences in diversity when stratified based on underlying IBD activity. These findings support our clinical finding that disease activity was not associated with an increased risk of CDI recurrence. Interestingly, when the microbiome of those with active IBD was analyzed, we noted a blunted increase in bacterial diversity in those that required medication escalation. Larger studies will be required to confirm this finding and to further assess the effect of medication initiation on the microbiome.

Our findings represent a single center experience and the retrospective nature of the study design is a limiting factor in data collection. Also, our definition of IBD flare relied on the determination of the treating physician. Furthermore, the number of patients in our microbiome analysis with active disease that required escalation is relatively small, although the differences in microbiome engraftment were significant after FMT and over time. The strengths of our study include the large number of subjects with and without IBD, allowing comparisons to be drawn between the two groups. Additionally, our cohort includes subjects with complex IBD and severe CDI, which are often excluded from other studies. The 6 month follow up period is also an important strength, as it provides a longer assessment of FMT efficacy relative to many studies.^6,7^ Lastly, the longitudinal microbiome analysis in a subset of our patients provides important results regarding microbial engraftment over time in relation to IBD activity and therapy.

In conclusion, our study shows FMT to be a successful treatment of recurrent or refractory CDI. Importantly, we did not find a difference in outcomes in subjects with or without IBD, supporting the hypothesis that underlying IBD does not decrease the efficacy of FMT. Microbiome analysis confirmed this observation, finding no difference between subjects with and without IBD. However, microbial engraftment was affected by IBD therapy, suggesting this is an important variable that should be accounted for in future studies.

## Supplementary Methods

### FMT procedure

FMT was performed preferentially via colonoscopy or flexible sigmoidoscopy following standard bowel lavage, with 250cc of FMT product instilled into the most proximal extent of exam. FMT performed via the upper gastrointestinal tract was performed via push enteroscopy, percutaneous endoscopic gastrostomy (PEG) tube, or jejunal (J) tube using 30cc of FMT product followed by a 30cc flush of non-bacteriostatic normal saline.

### Statistical analysis

R version 3.4.1 was utilized unless otherwise noted. The Log-rank test, presented as a Kaplan-Meier curve, was done utilizing R’s packages *survival* (version 3.1-131) and *survminer* (version 0.4.0). Changes in continuous outcomes over time were done using Linear Mixed-Effects Models using R’s package *NLME* (version 3.1-131) and *lsmeans* (version 2.27-2).

### Microbiome data generation and processing

Human fecal samples were collected fresh and stored at −80C prior to processing. Following suspension in extraction buffer, samples were mechanically lysed, centrifuged, and DNA extracted. The V4 variable region of the 16S rRNA gene was amplified by PCR using indexed primers as previously described^1^. Uniquely indexed 16S rDNA V4 amplicons were pooled and purified and the pooled samples were sequenced with an Illumina MiSeq (paired-end 250bp). Paired end reads were joined into a single DNA sequencing using the FLASH algorithm.^2^ We obtained a total of 7,263,850 reads (average 59,539 ± 34,744 reads/sample) after demultiplexing and quality filtering as previously described.^3^ Data was then clustered into Operational Taxonomic Units (OTUs) using a closed-reference OTU picking algorithm^4^ against Greengenes v13-8^5^, resulting in a total of 6,053 OTUs. Analysis of alpha and beta diversity was performed using QIIME v1.9.1^6^. Co-occurrence analysis of the microbiome immediately after FMT was performed using SparCC^7^ on the OTU table summarized at the genus level, and the resulting network was visualized using Cytoscape v3.0.2^8^ with network layout selected as edge-weighted Spring embedded metrics.

## Supplementary Tables

**SUPPLEMENTARY TABLE 1.**
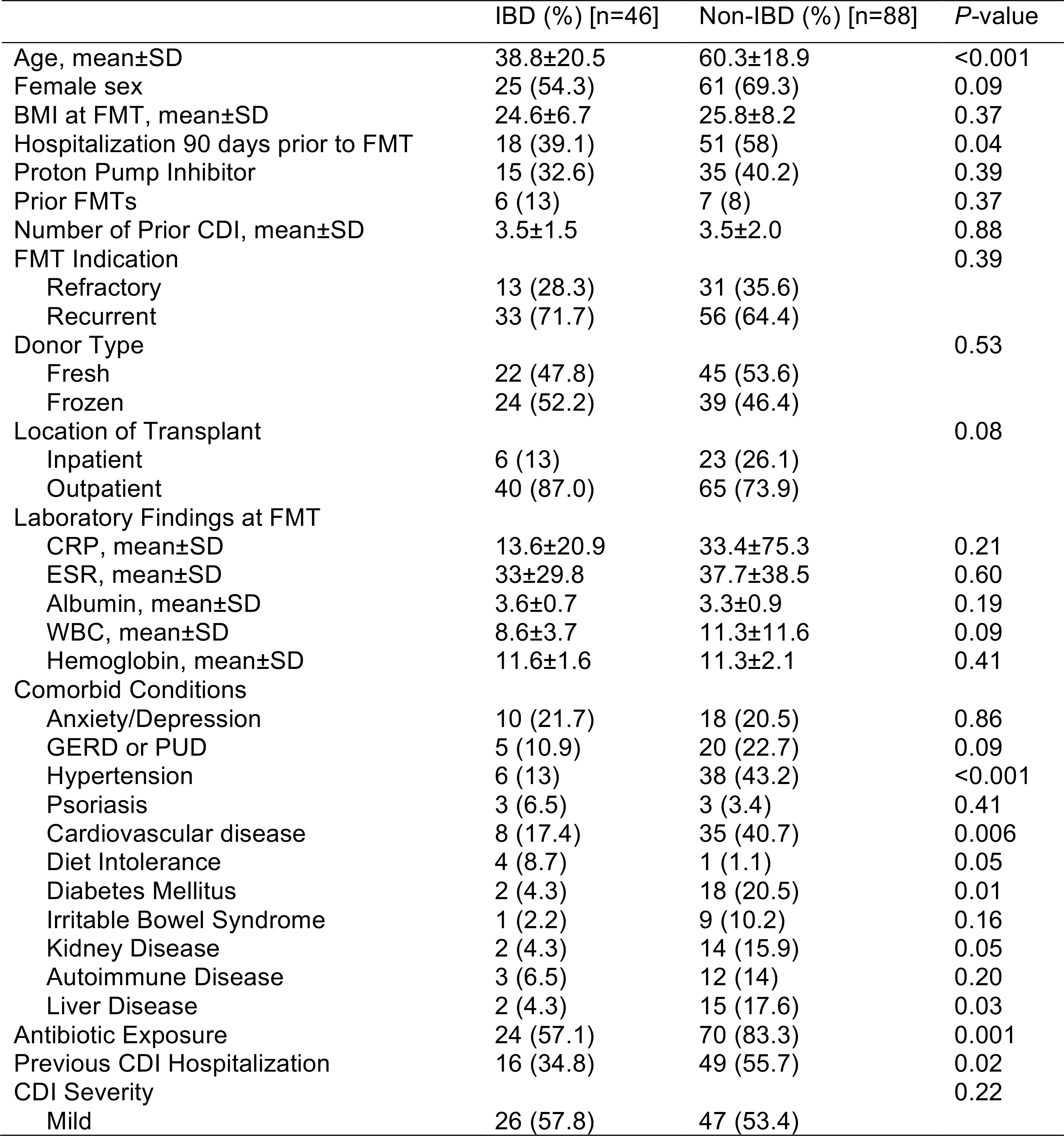
Patient Characteristics of the Cohort

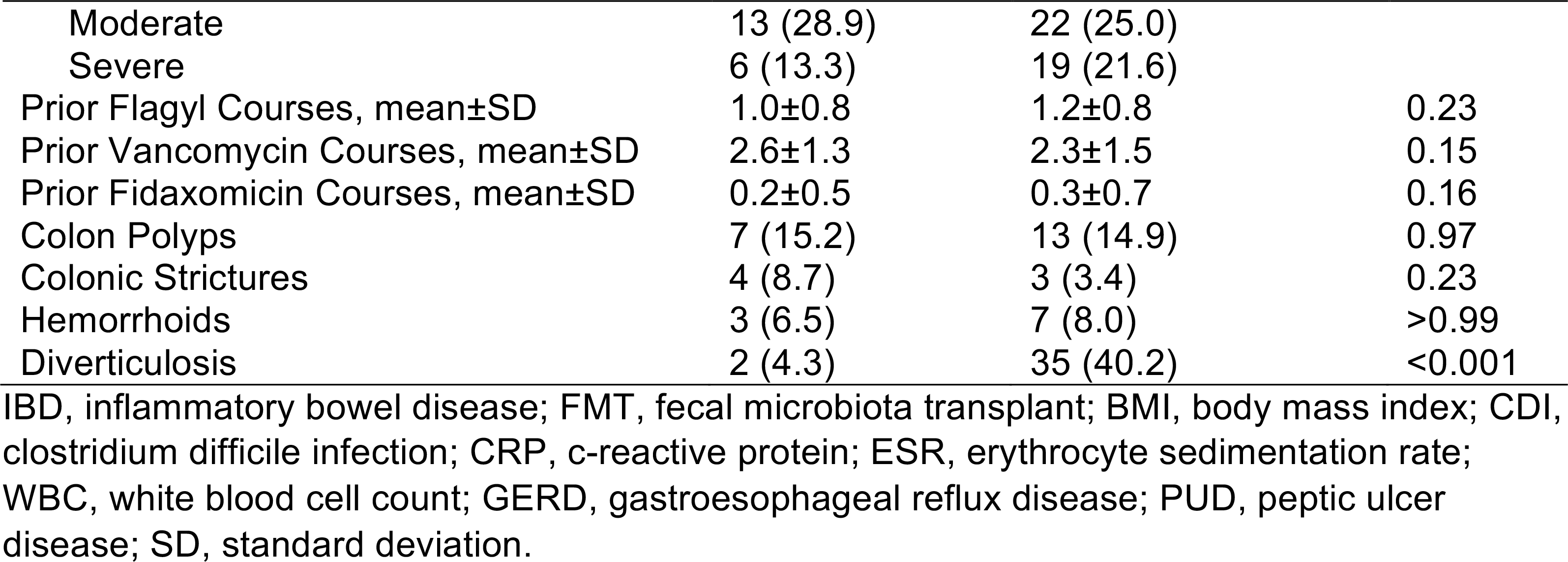

**SUPPLEMENTARY TABLE 2.** Characteristics of patients with inflammatory bowel disease at the time of FMT

|  | IBD (n=46) |
| --- | --- |
| IBD Type (%) |  |
| Crohn's Disease | 18 (39.1) |
| Ulcerative Colitis | 27 (58.7) |
| Indeterminate Colitis | 1 (2.2) |
| Endoscopic IBD Severity (%) |  |
| Remission | 8 (17.8) |
| Mild | 7 (15.6) |
| Moderate | 19 (42.4) |
| Severe | 11 (24.4) |
| Crohn's Disease Location (%) |  |
| Ileal | 1 (6.7) |
| Colonic | 8 (53.3) |
| Ileocolonic | 6 (40.0) |
| Ulcerative Colitis Location (%) |  |
| Proctitis | 2 (7.7) |
| Left Sided Colitis | 9 (34.6) |
| Extensive Colitis | 15 (57.7) |
| Indeterminate Colitis Location (%) |  |
| Colonic | 1 (100) |
| IBD Medication at the time of FMT (%)* |  |
| Mesalamine | 16 (34.8) |
| AZA/MP/MTX | 8 (17.4) |
| Steroids | 16 (34.8) |
| Anti-TNF | 21 (45.7) |
| Vedolizumab | 7 (15.2) |

IBD, inflammatory bowel disease; FMT, fecal microbiota transplant; AZA, azathioprine; MP, mercaptopurine; MTX, methotrexate; Anti-TNF, anti-tumor necrosis factor. \*Greater than 100% as patients are on combination therapy.

**SUPPLEMENTARY TABLE 3.**
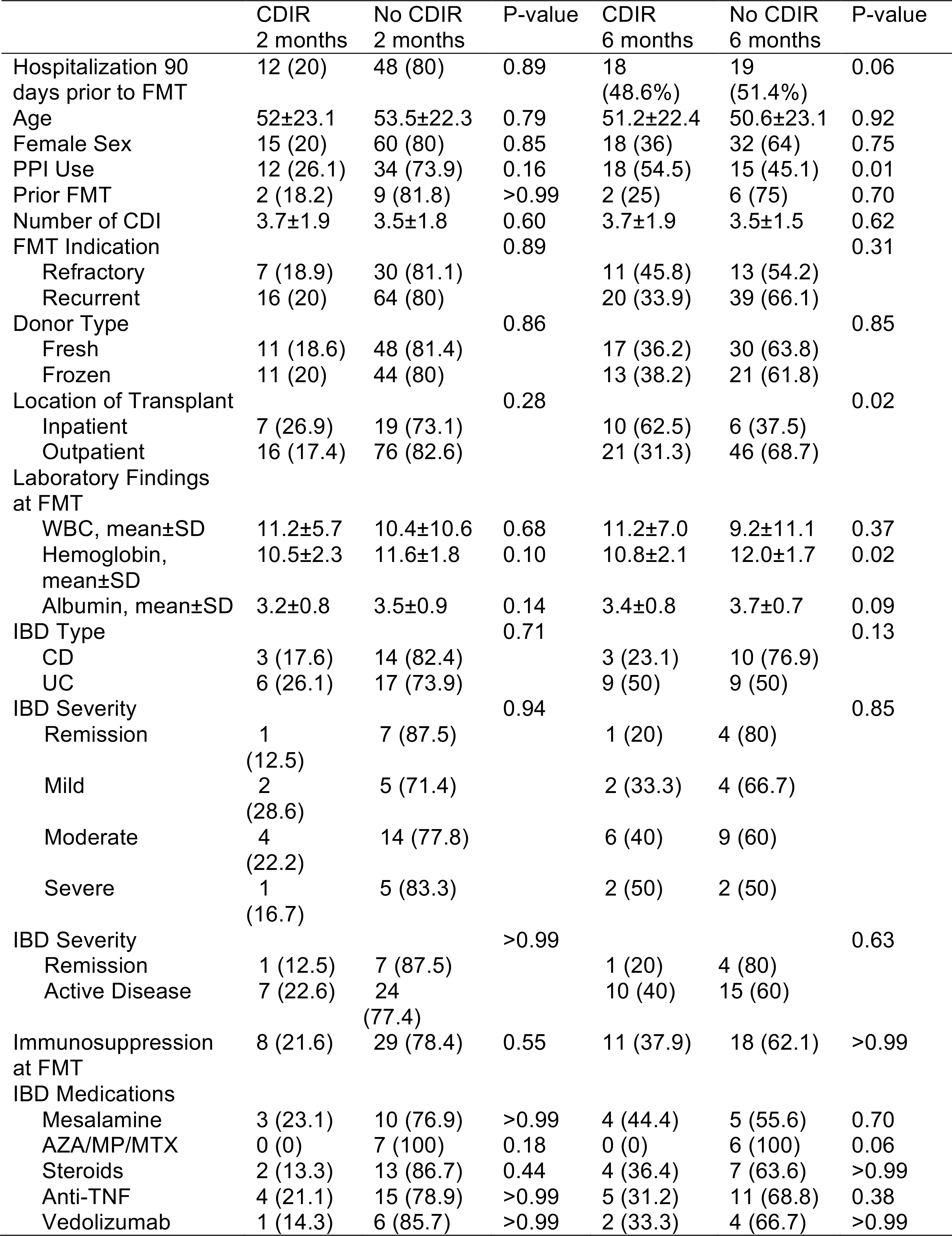
Univariate analysis for *Clostridium difficile* recurrence at 2 and 6 months

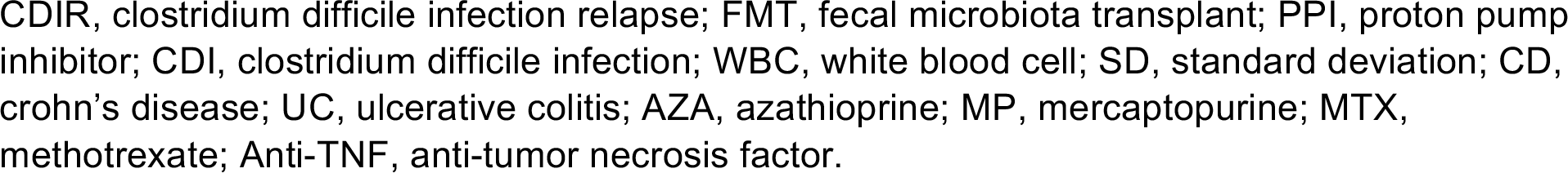

## Supplementary Figures

**SUPPLEMENTARY FIGURE 1.**
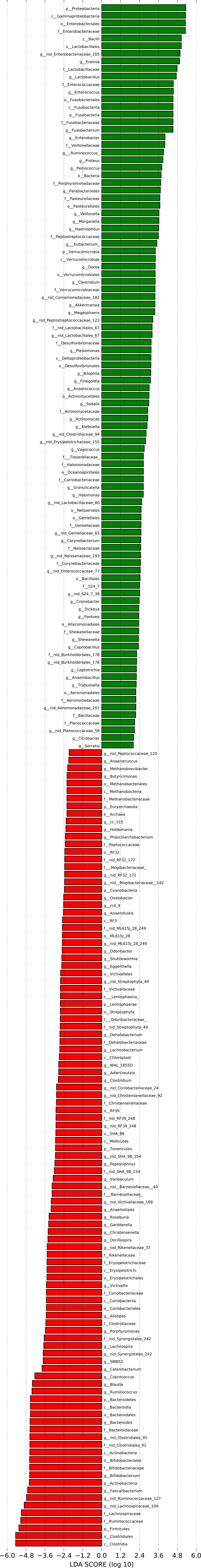
LEfSe analysis comparing microbial composition in donors (red) and patients (green) before FMT. Represented are all taxa significantly distinct with LDA scores > 2.0.

**SUPPLEMENTARY FIGURE 2.**
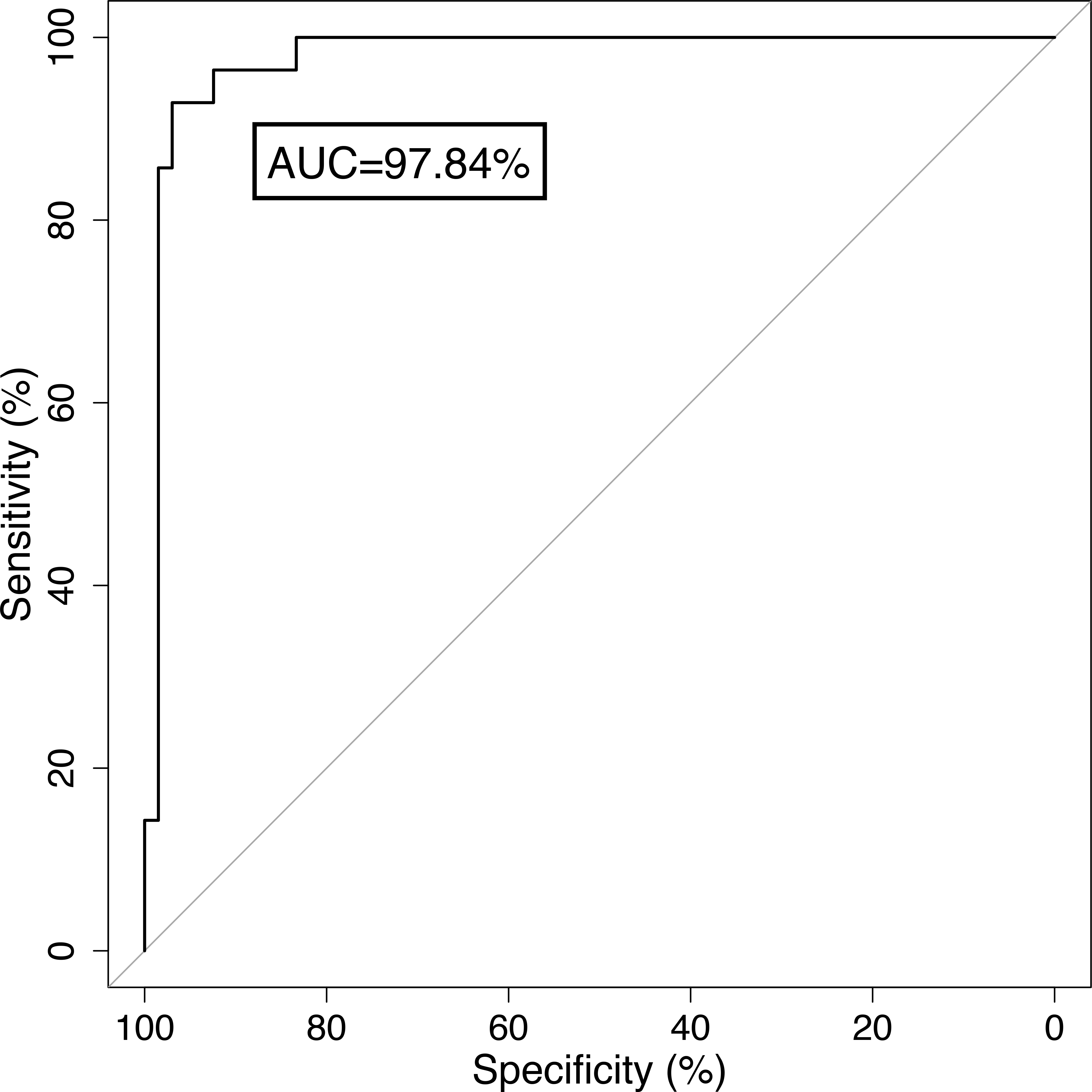
ROC curve for a random forest classifier trained to distinguish patients before and after FMT based on microbiome composition. AUC = area under the curve.

**SUPPLEMENTARY FIGURE 3.**
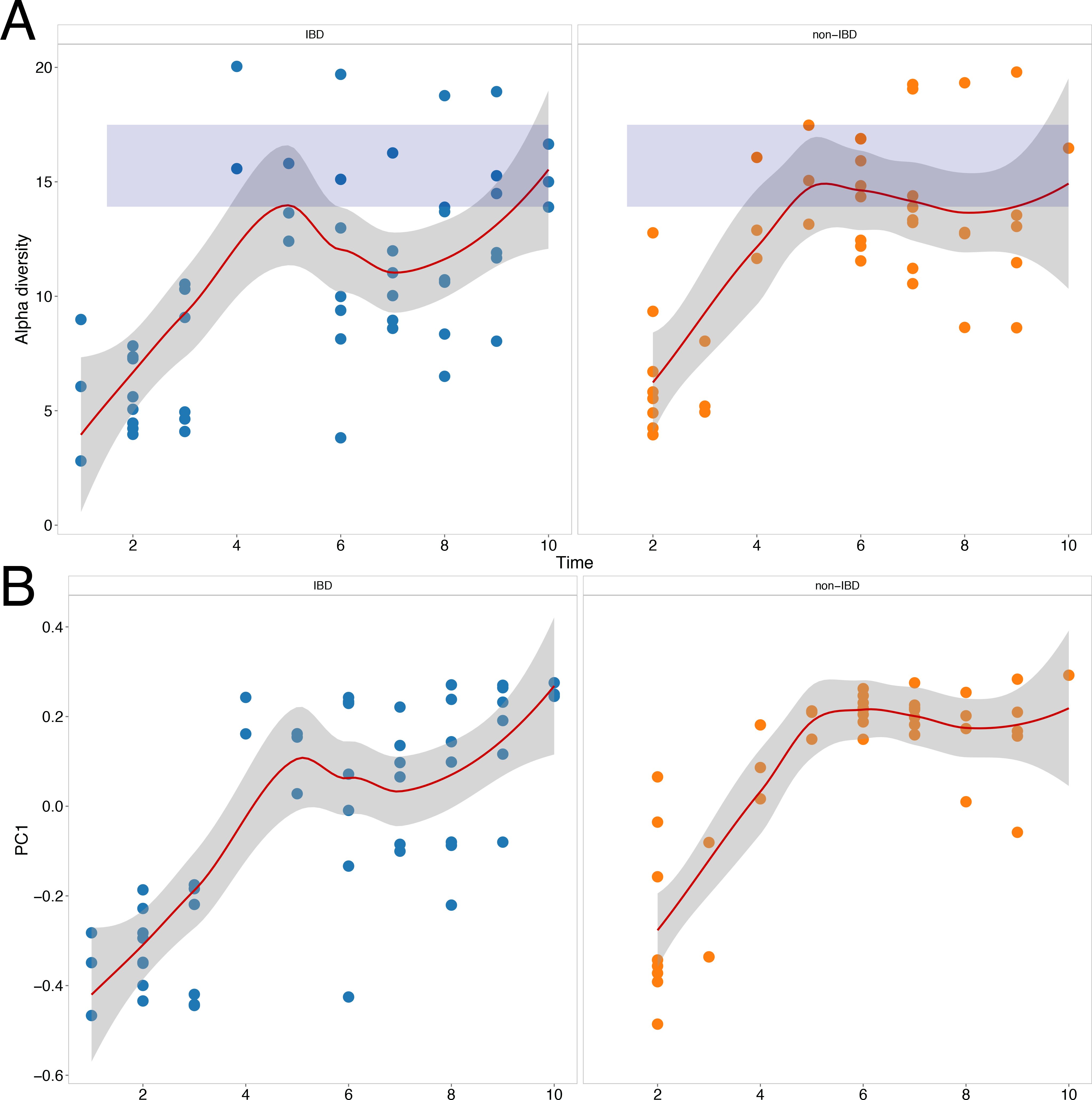
(A) Longitudinal alpha diversity over time in CDI patients with and without IBD. (B) First principal coordinate over time from PCoA analysis of beta diversity in CDI patients with and without IBD.

**SUPPLEMENTARY FIGURE 4.**
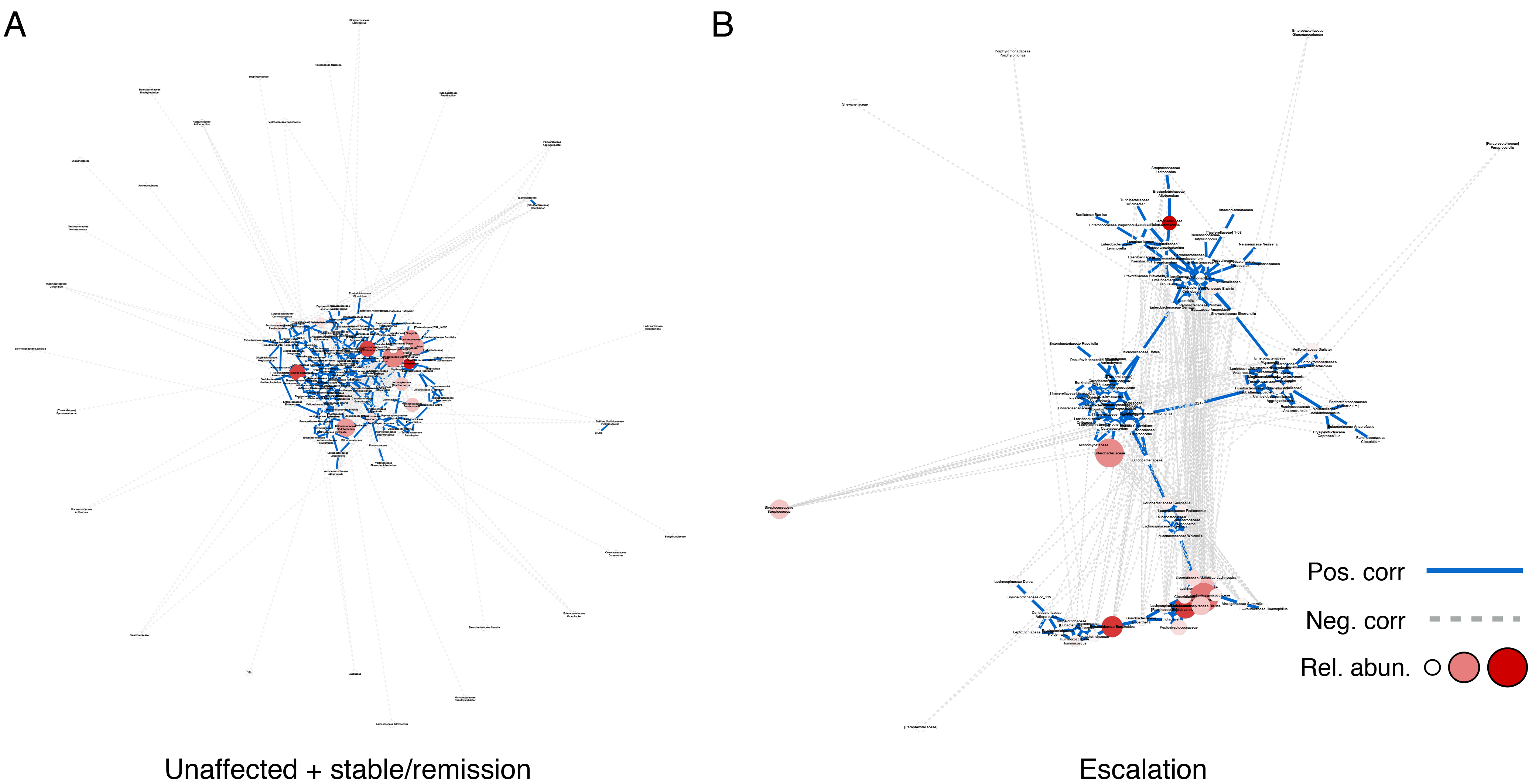
Co-occurrence network analysis of microbiome data after FMT in non-IBD/IBD^s^ (left) and IBD^e^ (right) patients. Nodes indicate bacterial taxa (summarized at the genus level), with size and color of the node being proportional to relative abundance of the taxa. Lines between nodes indicate correlations between taxa, with solid blue lines indicating positive correlations and dashed grey lines negative correlations.

Author Contributions: Conception and design: RPH, AG, SCF, JCC. Generation, collection, assembly, analysis, and/or interpretation of the data: RPH, AG, SCF, JCC, LY, MSF, JR, EJC, IM, NY, TL, PL, JJF. Drafting or revision of the manuscript: RPH, AG, SCF, LY, MSF, JR, EJC, IM, NY, TL, PRL, IP, JHC, BES, JFC, JJF, JCC. Approval of the final version of the manuscript: All authors read, reviewed, and approved the final version of the manuscript.

